# Evolution of BA.2.86 to JN.1 reveals functional changes in non-structural viral proteins are required for fitness of SARS-CoV-2

**DOI:** 10.1101/2025.02.17.638623

**Authors:** Shuhei Tsujino, Masumi Tsuda, Naganori Nao, Kaho Okumura, Lei Wang, Yoshitaka Oda, Yume Mimura, Jingshu Li, Rina Hashimoto, Yasufumi Matsumura, Rigel Suzuki, Saori Suzuki, Kumiko Yoshimatsu, Miki Nagao, The Genotype to Phenotype Japan (G2P-Japan) Consortium, Jumpei Ito, Kazuo Takayama, Kei Sato, Keita Matsuno, Tomokazu Tamura, Shinya Tanaka, Takasuke Fukuhara

**Affiliations:** Department of Microbiology and Immunology, Faculty of Medicine, Hokkaido University, Sapporo, Japan; Department of Cancer Pathology, Faculty of Medicine, Hokkaido University, Sapporo, Japan; Institute for Chemical Reaction Design and Discovery (WPI-ICReDD), Hokkaido University, Sapporo, Japan; One Health Research Center, Hokkaido University, Sapporo, Japan; Division of International Research Promotion, International Institute for Zoonosis Control, Hokkaido University, Sapporo, Japan; Institute for Vaccine Research and Development (IVReD), Hokkaido University, Sapporo, Japan; Division of Systems Virology, Department of Microbiology and Immunology, The Institute of Medical Science, The University of Tokyo, Tokyo, Japan; Faculty of Liberal Arts, Sophia University, Tokyo, Japan; Division of Risk Analysis and Management, International Institute for Zoonosis Control, Hokkaido University, Sapporo, Japan; Center for iPS Cell Research and Application (CiRA), Kyoto University, Kyoto, Japan; Department of Clinical Laboratory Medicine, Graduate School of Medicine, Kyoto University, Kyoto, Japan; Institute for Genetic Medicine, Hokkaido University, Sapporo, Japan; AMED-CREST, Japan Agency for Medical Research and Development (AMED), Tokyo, Japan; Graduate School of Frontier Sciences, The University of Tokyo, Chiba, Japan; Graduate School of Medicine, The University of Tokyo, Tokyo, Japan; International Research Center for Infectious Diseases, The Institute of Medical Science, The University of Tokyo, Tokyo, Japan; International Vaccine Design Center, The Institute of Medical Science, The University of Tokyo, Tokyo, Japan; Collaboration Unit for Infection, Joint Research Center for Human Retrovirus Infection, Kumamoto University, Kumamoto, Japan; MRC-University of Glasgow Centre for Virus Research, Glasgow, UK; International Collaboration Unit, International Institute for Zoonosis Control, Hokkaido University, Sapporo, Japan; Laboratory of Virus Control, Research Institute for Microbial Diseases, Osaka University, Suita, Japan; Department of Virology, Faculty of Medicine, Kyushu University, Fukuoka, Japan

**Author notes:** Correspondence (Tomokazu Tamura), (Shinya Tanaka), (Takasuke Fukuhara). These authors contributed equally (Shuhei Tsujino, Masumi Tsuda, Naganori Nao).

**Keywords:** SARS-CoV-2, COVID-19, JN.1, pathogenicity, recombinant virus, non-structural viral protein, S, NSP6, ORF7b

## Abstract

Severe acute respiratory syndrome coronavirus 2 (SARS-CoV-2), the causative agent of the coronavirus disease 2019 (COVID-19), is still circulating among humans, leading to the continuous evolution. SARS-CoV-2 Omicron JN.1 evolved from a distinct SARS-CoV-2 lineage, BA.2.86, spread rapidly worldwide. It is unclear why BA.2.86 did not become dominant and was quickly replaced by JN.1, which possesses one amino acid substitution in the spike protein (S:L455S) and two in the non-spike proteins NSP6 and ORF7b (NSP6:R252K and ORF7b:F19L) compared to BA.2.86. Here, we utilized recombinant viruses to elucidate the impact of these mutations on the virological characteristics of JN.1. We found that the mutation in the spike attenuated viral replication, but the non-spike mutations enhanced replication, suggesting the mutations in the non-spike proteins compensate for the one in the spike to improve viral fitness, as the mutations in the spike contribute to further immune evasion. Our findings suggest that functional changes in both the spike and non-spike proteins are necessary in the evolution of SARS-CoV-2 to enable evasion of adaptive immunity within the human population while sustaining replication.

**IMPORTANCE:** Because the spike protein is strongly associated with certain virological properties of SARS-CoV-2, such as immune evasion and infectivity, most previous studies on SARS-CoV-2 variants have focused on spike protein mutations. However, the non-spike proteins also contribute to infectivity, as observed throughout the evolution of Omicron subvariants. In this study, we demonstrate a “trade-off” strategy in SARS-CoV-2 Omicron JN.1 in which the reduced infectivity caused by spike mutation is compensated by non-spike mutations. Our results provide insight into the evolutionary scenario of the emerging virus in the human population.

## INTRODUCTION

Since its emergence in 2019, SARS-CoV-2, the causative agent of coronavirus disease 2019 (COVID-19), has led to a global pandemic. Early SARS-CoV-2 evolved toward increased pathogenicity, but due to its spread and increased human vaccination, Omicron subvariants have diminished pathogenicity and enhanced immune escape compared with ancestral variants (1–10). Importantly, these subvariants continue to circulate in human populations.

Two dramatic events have occurred in the evolution of SARS-CoV-2 during the pandemic. The first was the emergence of Omicron (BA.1) from the Delta subvariant, and the second was the evolution from Omicron BA.2 to BA.2.86 (2, 10). Between Delta and BA.1, 38 mutations were identified in the viral spike (S) protein and 49 in other viral genes. Similarly, BA.2.86 exhibited 32 mutations in S and 14 in other genes compared to BA.2 (Nextstrain; https://nextstrain.org/ncov/gisaid/global/6m) (2, 10). Then, just as BA.2 emerged from BA.1, the descendant JN.1 emerged in the United States in September 2023 and outcompeted BA.2.86 to become the dominant variant (11). As of December 2024, direct descendants of JN.1, including KP.3 and KP.3.1.1, have become predominant globally (12, 13).

Because the JN.1 lineage surged and rapidly outcompeted previously dominant variants in early 2024, the effective reproduction number (R_e_) and immune-evasive properties of the JN.1 variant have been of great interest to researchers, including ourselves (11, 14, 15). JN.1 showed even greater immune evasion than BA.2.86 but exhibited reduced binding affinity for the SARS-CoV-2 receptor angiotensin-converting enzyme 2 (ACE2). Cryo-EM observations revealed that the mutation L455S in the receptor-binding domain (RBD) of the JN.1 S protein disrupted the interaction between RBD and human ACE2 (16). These findings suggest that the lower affinity of the JN.1 vs BA.2.86 S protein for ACE2 impairs viral entry and viral adaptation against host immune defenses. Thus, it remains an open question how JN.1 outcompeted BA.2.86 to quickly become dominant.

Reverse genetics systems have played a central role in studying viral replication, pathogenicity, and the impact of specific mutations. Since a rapid reverse genetics system for SARS-CoV-2 has been developed (17, 18), we have investigated the viral characteristics of Delta, BA.2, XBB.1.5, and EG.5.1, elucidating the roles of specific mutations (1, 8, 9, 19). Here, in this study, we used recombinant viruses to examine the differences in viral characteristics between JN.1 and its direct ancestor, BA.2.86. To this end, we analyzed the mutation frequency and identified the mutation(s) that characterize JN.1. Furthermore, we generated recombinant viruses with these mutations and investigated their replication efficiency and intrinsic pathogenicity.

## RESULTS AND DISCUSSION

To investigate the replication efficiency and intrinsic pathogenicity, we inoculated VeroE6 cells expressing transmembrane serine protease 2 (TMPRSS2) (20) with clinical isolates of JN.1 (GISAID ID: EPI_ISL_18771637) or BA.2.86 (GISAID ID: EPI_ISL_18233521) (10). In quantifying the infectious viral titers of the supernatants, we found that the replication properties of BA.2.86 and JN.1 were comparable (Fig. 1A). Next, we intranasally inoculated hamsters―our established animal model for COVID-19 (1–10)―with either BA.2.86 or JN.1 under anesthesia. The body weights of the hamsters were comparable for the two viruses and, as expected, lower than those of uninfected hamsters (Fig. 1B). These findings suggest that under our experimental conditions, the replication efficiency and intrinsic pathogenicity of JN.1 and BA.2.86 are similar.

**Fig. 1.**
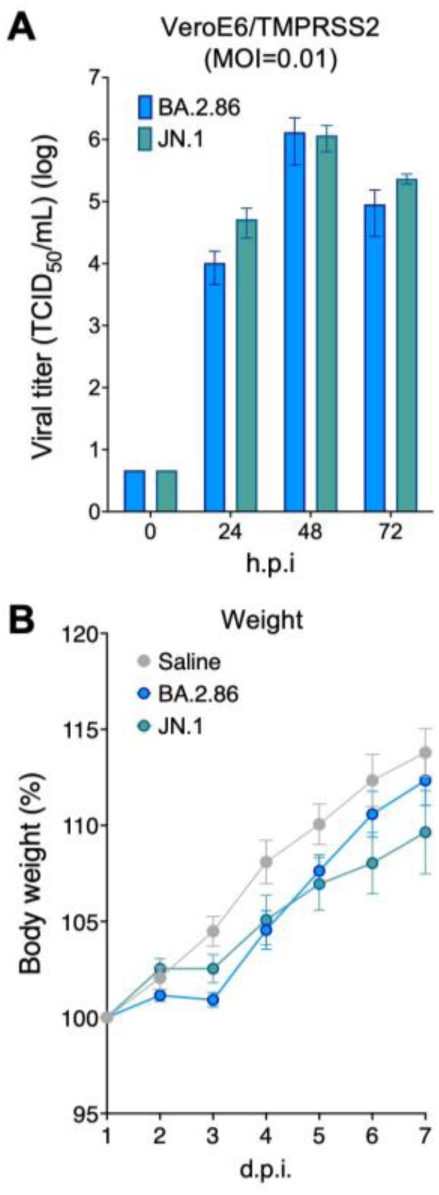
Virological characteristics of SARS-CoV-2 BA.2.86 and JN.1. **(A)** BA.2.86 and JN.1 were used to inoculate VeroE6/TMPRSS2 cells (MOI = 0.01). The 50% tissue culture infectious dose (TCID_50_) of the culture supernatants were quantified at the indicated times post infection. **(B)** Syrian hamsters were intranasally inoculated with clinical isolates of BA.2.86 or JN.1 (5,000 TCID_50_) or, as a negative control, saline (each n=6). Data are represented as mean ± SEM. Body weight was tracked daily through 7 d.p.i. h.p.i.: hours post-infection; d.p.i.: days post-infection.

To investigate the evolution of BA.2.86 to JN.1, the frequency of mutations across multiple Omicron subvariants was examined (Fig. 2A). In comparing BA.2.86 to JN.1, two substitutions in non-spike proteins (NSP6 and ORF7b) were identified in a minor population of BA.2.86. An additional amino acid substitution in S (S:L455S) was also acquired and, then, the three mutations (S:L455S, NSP6:R252K and ORF7b:F19L) that have been documented as convergent substitutions (https://jbloomlab.github.io/SARS2-d-fitness/; Access date: August 03, 2024). The JN.1 variant was first detected in September 2023, and became globally predominant by January 2024 (Nextstrain, clade 24A; https://nextstrain.org/ncov/gisaid/global/6m) (11). It subsequently diverged into several descendant lineages, including KP.2, KP.3 and KP.3.1.1 (Pango nomenclature; https://github.com/cov-lineages/pango-designation) (12, 13, 21). Of note, the three mutations that initially emerged in JN.1 remain present in currently circulating variants such as KP.3.1.1 (Fig. 2A).

**Fig. 2.**
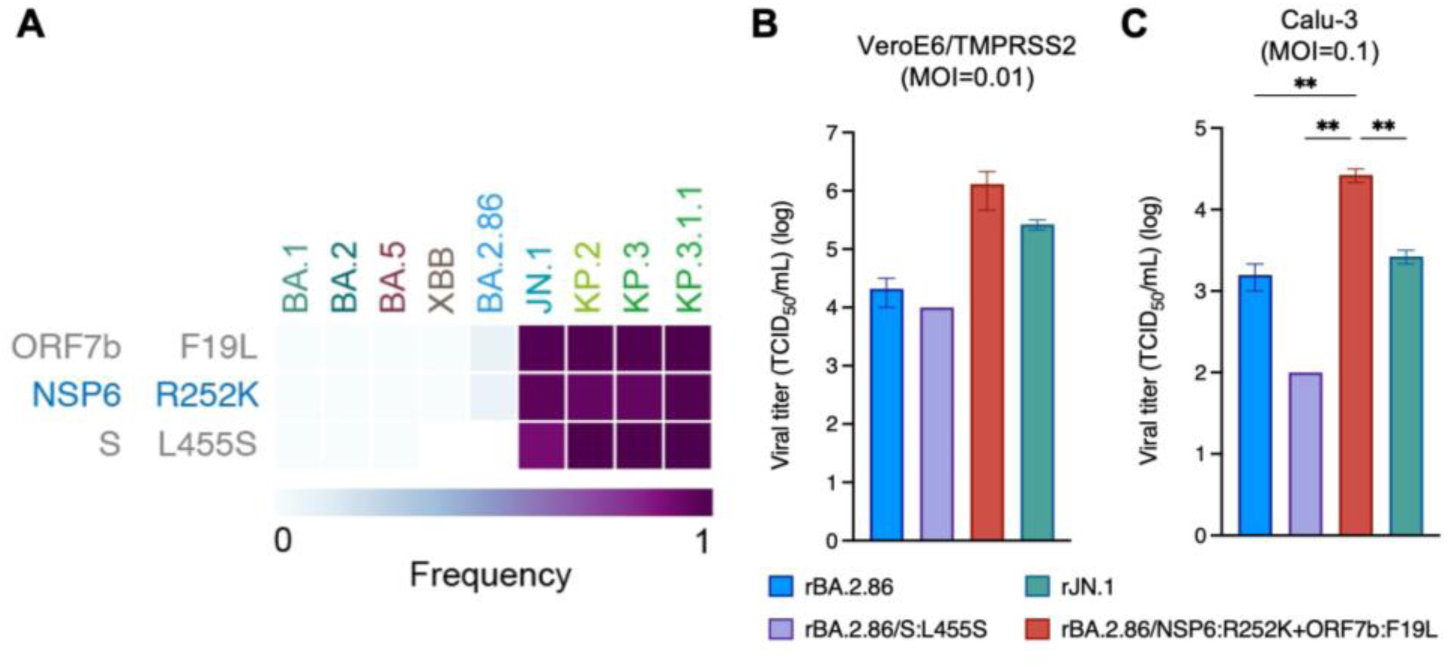
Spike and non-spike mutations define the evolution of BA.2.86 to JN.1. **(A)** Frequency of mutations in BA.1, BA.2, BA.5, XBB, BA.2.86, JN.1, KP.2, KP.3, and KP.3.1.1. Only mutations that differ between JN.1 and its parental lineage, BA.2.86, are shown. **(B and C)** Growth of rBA.2.86, rJN.1, rBA.2.86/S:L455S, and rBA.2.86/NSP6:R252K+ORF7b:F19L in the supernatants of VeroE6/TMPRSS2 cells (MOI = 0.01) **(B)** and Calu-3 cells (MOI = 0.1) **(C)** at 24 h.p.i. Data are represented as mean ± SEM.

Previous studies, including ours, showed that the S:L455S mutation in JN.1 has a negative effect on RBD-ACE2 binding affinity (11, 16). However, our *in vitro* analysis demonstrated similar growth kinetics for BA.2.86 and JN.1 (Fig. 1A). To further evaluate the impact of spike and non-spike mutations on viral growth in cell culture, we generated recombinant viruses: rBA.2.86; rJN.1; rBA.2.86/NSP6:R252K+ORF7b:F19L (a mutant carrying the two substitutions in NSP6 and ORF7b described above that initially emerged in humans); and rBA.2.86/S:L455S (a mutant carrying the S:L455S mutation).

Calu-3 and VeroE6/TMPRSS2 cells were inoculated with each of these four recombinant viruses. In both cell types, growth of rBA.2.86 and rJN.1 were almost identical (Fig. 2B and C), consistent with our results using the clinical isolates. In the supernatants of VeroE6/TMPRSS2 cells, the titers of rBA.2.86/NSP6:R252K+ORF7b:F19L were higher than those of rBA.2.86 (Fig. 2B). In Calu-3 cells, the replication kinetics of rBA.2.86/S:L455S were lower than those of rBA.2.86. On the other hand, the growth of rBA.2.86/NSP6:R252K+ORF7b:F19L were significantly higher than those of rBA.2.86 (Fig. 2C). These results suggest that the S:L455S mutation decreases replication efficiency, while the NSP6:R252K and ORF7b:F19L mutations increase replication efficiency *in vitro*.

NSP6, a multi-spanning transmembrane protein, is involved in the biogenesis of the viral replication complex (22). The ORF7b protein has been reported to facilitate viral infection and production and to inhibit the RIG-I-like receptor (RLR) signaling pathway (23). The mutations in these non-spike proteins in JN.1 may affect replication efficiency. To further investigate this possibility, two additional recombinant JN.1 viruses containing the mutations to revert to the BA.2.86 sequence were generated (rJN.1/NSP6:K252R and rJN.1/ORF7b:L19F). rJN.1 containing an S mutation to revert back to the BA.2.86 sequence was also evaluated. While designated in these experiments as rJN.1/S:S455L, this virus is the same sequence as rBA.2.86/NSP6:R252K+ORF7b:F19L.

In VeroE6/TMPRSS2 cells, the growth of rJN.1/S:S455L was significantly higher than those of rBA.2.86, consistent with the greater cytopathic effect observed (Fig. 3A and C). In Calu-3 cells (Fig. 3B), the growth of rJN.1/S:S455L were significantly higher than those of rBA.2.86 and rJN.1. In both cell lines, the non-spike protein mutants (rJN.1/NSP6:K252R and rJN.1/ORF7b:L19F) did not significantly affect growth compared to the parental rJN.1. Altogether, these results in Figure 2 suggest that in evolving from BA.2.86 to JN.1, the S:L455S mutation attenuates replication efficiency *in vitro*, while mutations in NSP6 and ORF7b contribute to higher replication efficiency.

**Fig. 3.**
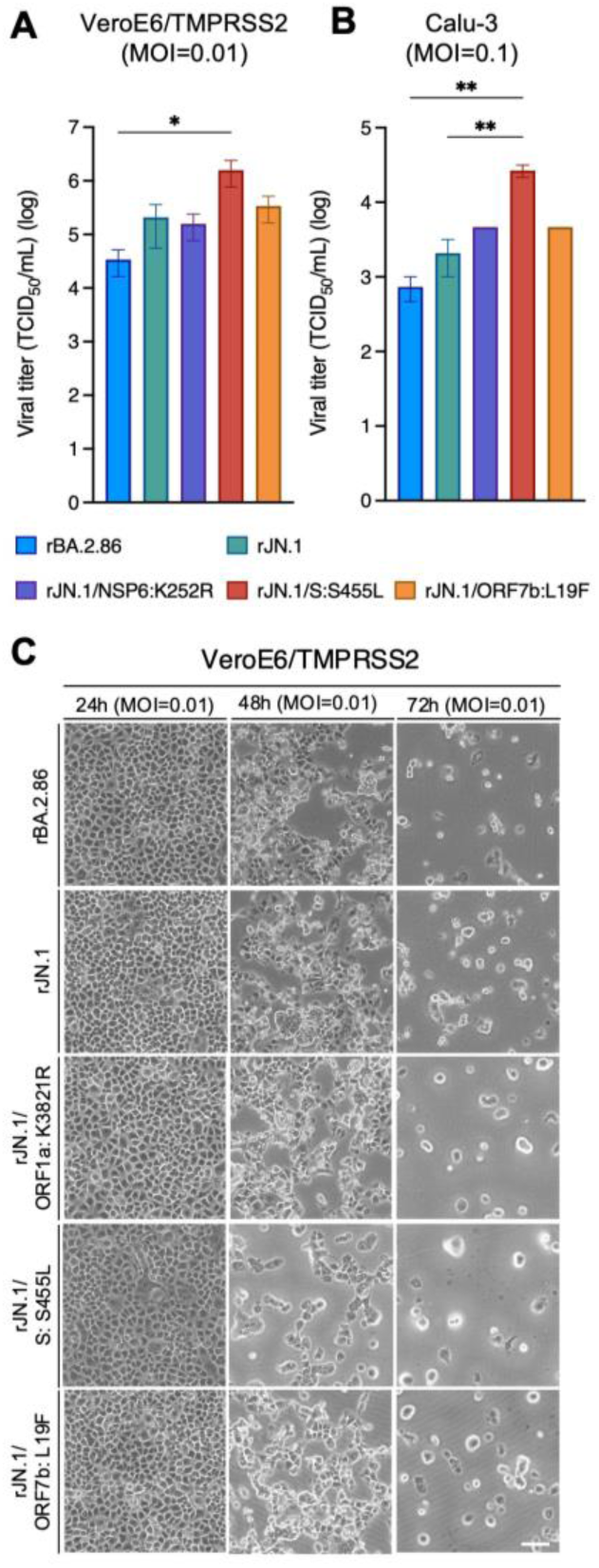
The impact of single mutations on the virological characteristics of JN.1. **(A and B)** Growth of rBA.2.86, rJN.1, rJN.1/S:S455L, rJN.1/NSP6:K252R, and rJN.1/ORF7b:L19F in the supernatants of VeroE6/TMPRSS2 cells (MOI = 0.01) **(A)** and Calu-3 cells (MOI = 0.1) **(B)** at 24 h.p.i. **(C)** The VeroE6/TMPRSS2 cells were examined by bright field microscopy at the indicated times post infection to assess cytopathic effect. Data are represented as mean ± SEM. Scale bars, 500 μm.

To investigate the *in vivo* dynamics and pathogenicity of these viruses, Syrian hamsters were intranasally inoculated with rBA.2.86, rJN.1, and the different rJN.1 mutants. Consistent with the in vitro findings for the clinical isolates, changes in weight were comparable between hamsters infected with rJN.1 vs rBA.2.86. Of the two single-mutation rJN.1 viruses, only rJN.1/S:S455L compared to rJN.1 led to significant weight loss, which was also the greatest among all viruses evaluated (Fig. 4A, left). On the other hand, the weight loss of hamsters infected with rJN.1/NSP6:K252R were significantly lower than those of hamsters infected with rJN.1. Hamsters infected with rJN.1/ORF7b:L19F only showed slightly less weight loss compared to hamsters infected with rJN.1.

**Fig. 4.**
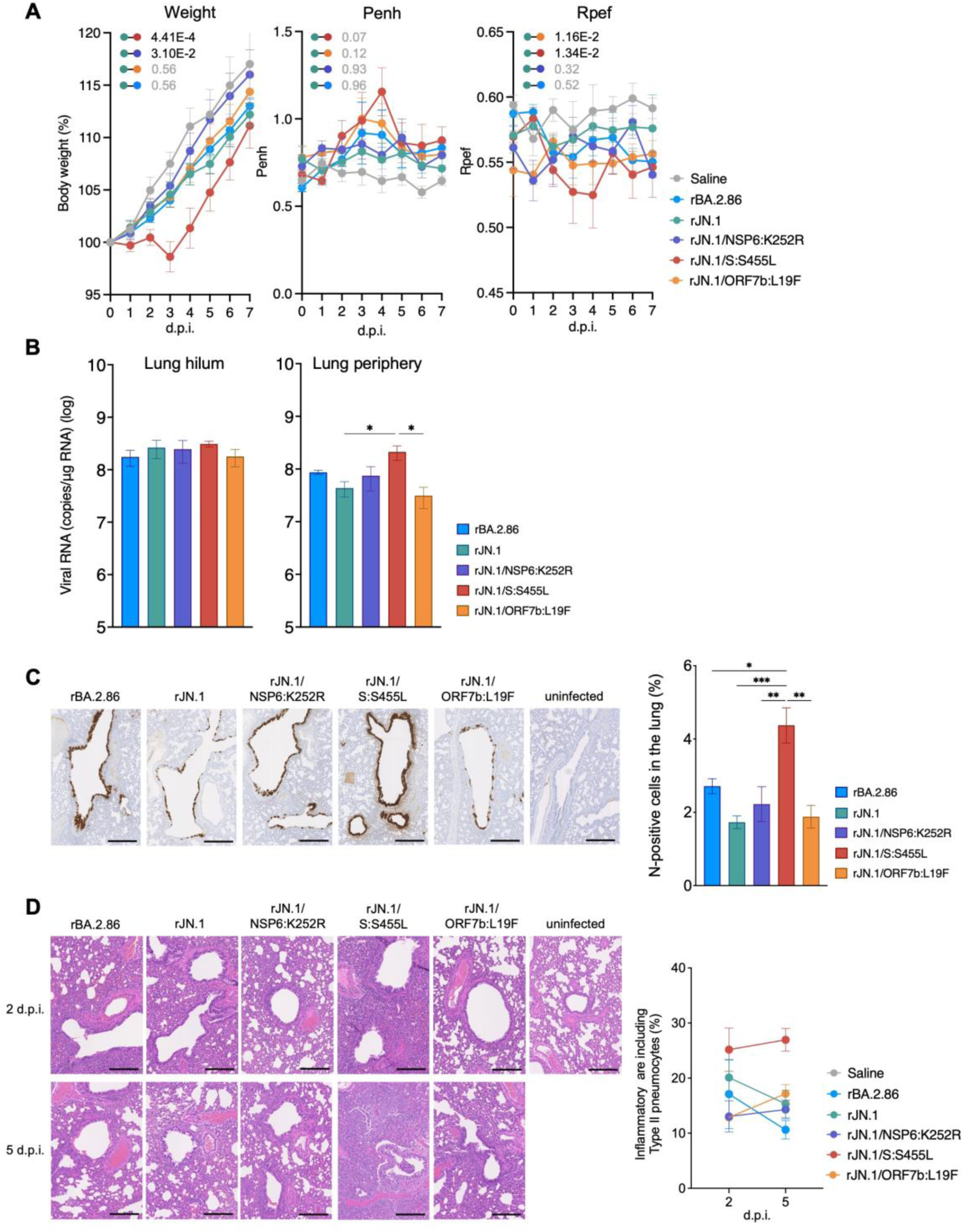
Mutations in non-spike proteins underlie the immunopathogenic features of JN.1. Syrian hamsters were intranasally inoculated with JN.1 backbone viruses (5,000 TCID_50_) or saline (uninfected). **(A)** Body weight (left), enhanced pause (Penh; middle), and ratio of time to peak expiratory flow relative to the total expiratory time (Rpef; right) of infected hamsters (n=6 hamster of the same age per infection/control group). The familywise error rates calculated using the Holm method are indicated in the figures. **(B)** Viral RNA loads in the lung hilum (left) and lung periphery (right) of infected hamsters (n = 4 per infection group) at 2 d.p.i. **(C)** Immunohistochemistry of the viral N protein (brown staining) in hamster lung tissue at 2 d.p.i. Representative figures are shown. Scale bars, 500 µm. **(D)** Hematoxylin and eosin staining of the lungs at 2 d.p.i. (upper) and 5 d.p.i. (lower) of infected/control hamsters. Representative figures are shown. Data are represented as mean ± SEM. Scale bars, 250 µm.

SARS-CoV-2 infection causes a decline in pulmonary function (24), and the degree of deterioration can be used as an index of viral pathogenicity (2). Thus, we analyzed the pulmonary function of infected hamsters using two parameters, enhanced pause (Penh) and the ratio of time to peak expiratory flow relative to the total expiratory time (Rpef). Of the viruses evaluated, both rJN.1/S:S455L and rJN.1/ORF7b:L19F significantly impaired Rpef compared to rJN.1 (Fig. 4A, middle and right). rJN.1/S:S455L infection also resulted in significantly higher Penh compared to compared to rJN.1 and was the only one of the viruses evaluated to do so.

Moreover, to evaluate viral spread in respiratory tissues, we collected the lungs of infected hamsters at 2- and 5-days post-infection (d.p.i.), separating the tissues into the hilum and periphery regions. In the hilum, the viral RNA load of rJN.1/S:S455L-infected hamsters was comparable to that of rJN.1-infected hamsters (Fig. 4B, left). In contrast, in the lung periphery region, the viral RNA loads of the rJN.1/S:S455L-infected hamsters was significantly higher than those of the rJN.1- and rJN.1/ORF7b:L19F-infected hamsters (Fig. 4B, right). These results suggest that the efficacy of viral spread in the lung is greater with rJN.1/S:S455L vs rJN.1 or rJN.1/ORF7b:L19F.

We also performed immunohistochemistry (IHC) to evaluate the presence of viral N protein in the respiratory tissues of infected hamsters. In the lung 2 d.p.i., N-positive cells were more strongly detected in the bronchial/bronchiolar epithelia of rJN.1/S:S455L-infected hamsters than rBA.2.86-, rJN.1-, rJN.1/NSP6:K252R- or rJN.1/ORF7b:L19F-infected hamsters (Fig. 4C). Then, to evaluate severity of inflammation upon infection with the mutant viruses, histopathological analyses were performed on the lung tissue. At 2 d.p.i., alveolar damage around the bronchi was prominent in rJN.1/S:S455L-infected hamsters (Fig. 4D). On the other hand, inflammation in bronchi/bronchioles tended to be more limited in rJN.1/NSP6:K252R- and rJN.1/ORF7b:L19F-infected hamsters than in rBA.2.86 and rJN.1. At 5 d.p.i., the alveolar architecture appeared more severely destroyed by alveolar damage or the expansion of type II pneumocytes in rJN.1/S:S455L-infected hamsters (Fig. 4D). No significant differences were found between rBA.2.86-, rJN.1-, rJN.1/NSP6:K252R- and rJN.1/ORF7b:L19F-infected hamsters. Taken together, these findings suggest that the S:L455S mutation acquired in the evolution of BA.2.86 to JN.1 attenuates viral growth and pathogenicity.

This study aimed to investigate the virological characteristics of SARS-CoV-2 JN.1 and understand the evolution of this variant from BA.2.86. Our findings suggest that the balance between properties of the spike and non-spike proteins is important for viral fitness and continued circulation of SARS-CoV-2 in humans. Compared with its direct ancestor BA.2.86, JN.1 has three mutations: S:L455S, NSP6:R252K, and ORF7b:F19L (Fig. 2A). Using recombinant mutant viruses generated in both the BA.2.86 and JN.1 backbones, we demonstrated that the S:L455S mutation attenuates replication efficiency and pathogenicity, while the NSP6:R252K and ORF7b:F19L mutations appear to compensate for this attenuation. BA.2.86 did not become predominant in human populations and was quickly replaced by JN.1 (25). This is likely because BA.2.86 exhibited weaker immune evasion than previously dominant variants (26–28). Acquisition of the S:L455S mutation may have helped enhance immune evasion but at the cost of impaired viral replication. Thus, additional mutations in non-spike proteins (NSP6:R252K and ORF7b:F19L), which had already been observed in a minor population of BA.2.86, increased the viral fitness of JN.1 and enabled more efficient circulation in humans.

Throughout the evolution of Omicron subvariants, SARS-CoV-2 has demonstrated improved immune evasion while maintaining infectivity (29). Acquisition of mutations in the spike protein imposes a weakness on viral fitness. To overcome this, mutations in non-spike proteins could enhance viral fitness by modulating certain virological properties. This “trade-off” strategy has been consistently observed during the circulation of Omicron subvariants in humans. For instance, in BA.1, mutations in both spike and NSP6 were reported to contribute to attenuated pathogenicity (30). In BA.2, we reported that the spike mutation S:L371F enhanced fusogenicity and pathogenicity while multiple non-spike mutations attenuated replication efficiency and pathogenicity (19). Furthermore, the impairment of major histocompatibility complex suppression due to dysfunctional ORF8 in XBB.1.5 was shown to influence viral pathogenicity (8). All this knowledge taken together demonstrates that investigations into the impact of mutations not only in the spike protein but also in the non-spike proteins are crucial for understanding SARS-CoV-2 evolution.

As we demonstrated for a variety of SARS-CoV-2 Omicron subvariants in the past (2–13, 19, 21), elucidating the virological features of newly emerging SARS-CoV-2 variants is important to determine their potential risk to human society and to understand the evolution of this virus in humans. Accumulating knowledge of the evolutionary traits of newly emerging pathogenic viruses in the human population will be beneficial in preparing for future outbreaks and pandemics.

## MATERIALS AND METHODS

### Ethics statement

All experiments with hamsters were performed in accordance with the Science Council of Japan’s Guidelines for the Proper Conduct of Animal Experiments. The protocols were approved by the Institutional Animal Care and Use Committee of National University Corporation Hokkaido University (approval ID: 20-0123).

### Cell culture

VeroE6/TMPRSS2 cells (VeroE6 cells stably expressing human TMPRSS2; JCRB Cell Bank, JCRB1819) (20) were maintained in DMEM (low glucose) (Cat#041-29775; FUJIFILM WAKO, Osaka, Japan) containing 10% FBS, 1 mg/mL G418 (Cat#09380-44; Nacalai Tesque, Kyoto, Japan). Calu-3 cells (ATCC, HTB-55) were maintained in Eagle’s minimum essential medium (EMEM) (Cat#056-0838; Sigma-Aldrich, MO, USA) containing 10% FBS and 1% penicillin-streptomycin (Cat#09367-34; Nacalai Tesque).

### Epidemic dynamics analysis and mutation frequency calculations

In this study, we analyzed the viral genomic surveillance data stored in the GISAID database (https://www.gisaid.org; downloaded on September 11, 2024) (31). We used the data collected for SARS-CoV-2 from April 1, 2022 to September 1, 2024 in this analysis. We excluded any data that i) did not have a collection date and Pango lineage information; ii) was retrieved from non-human animals; and iii) was sampled during quarantine. Only BA.1, BA.2, BA.5, XBB, BA.2.86, JN.1, KP.2, KP.3, and KP.3.1.1 variants are included in Figure 2A. Mutation frequency of each lineage was calculated by dividing the number of sequences harboring the substitution of interest with the total number of sequences in each lineage.

### Plasmid construction

The nine pmW118 plasmids containing the partial genomes of SARS-CoV-2 BA.2.86 were previously generated (32). To generate the recombinant JN.1 viruses, mutations were introduced by inverse fusion PCR cloning into the plasmids encoding the corresponding BA.2.86 genes. Sequences of all the plasmids used in this study were confirmed by a SeqStudio Genetic Analyzer (Thermo Fisher Scientific, MA, USA) and an outsourced service (Fasmac, Kanagawa, Japan). Primer and plasmid information can be provided upon request.

### SARS-CoV-2 preparation and titration

The working stocks of SARS-CoV-2 virus were prepared and titrated as previously described (8). In this study, stocks were prepared using clinical isolates of BA.2.86 (strain TKYnat15020; GISAID ID: EPI_ISL_18233521) (10) and JN.1 (strain LG0688; GISAID ID: EPI_ISL_18771637).

Recombinant viruses were generated by a circular polymerase extension reaction (CPER) (17). The resultant CPER products were transfected into VeroE6/TMPRSS2 cells as described previously (8). All the viruses were stored at −80°C until use and viral genome sequences were confirmed by Sanger sequencing (see “Plasmid construction” section above).

### Titration and growth kinetics

The infectious titers of supernatants from infected cell cultures were determined by quantifying the 50% tissue culture infectious dose (TCID_50_) (33). For growth kinetics, VeroE6/TMPRSS2 cells or Calu-3 cells were inoculated with virus in 12-well plates at a multiplicity of infection (MOI) of 0.01 or 0.1, respectively. The infectious titers of supernatants collected at the indicated timepoints were then determined.

### Assessment of viral pathogenicity in hamsters

Animal experiments were performed as previously described (1–10). In brief, Syrian hamsters (males, 4 weeks old) were intranasally inoculated under anesthesia with virus (5,000 TCID_50_ in 100 μL) or saline (100 μL). Body weight was recorded daily until 7 d.p.i. Enhanced pause (Penh) and the ratio of time to peak expiratory follow relative to the total expiratory time (Rpef) were measured using a Buxco Small Animal Whole Body Plethysmography system (Data Sciences International, MN, USA) every day until 7 d.p.i. Lung tissues were collected at 2 and 5 d.p.i. The viral RNA load in the respiratory tissues was determined by RT-qPCR using a QuantStudio 5 Real-Time PCR system (Thermo Fisher Scientific), as described previously (34, 35). These tissues were also used for immunohistochemistry and hematoxylin and eosin staining as previously described (1–10, 19). Expression of viral proteins was visualized using anti-SARS-CoV-2 N monoclonal antibody (clone 1035111, R&D Systems, 1:400). Images were incorporated as virtual slides by NDP.scan software v3.2.4 (Hamamatsu Photonics, Shizuoka, Japan). The area of N-protein positivity and inflammation was measured using Fiji software v2.2.0 (ImageJ) according to the criteria of certified pathologists (1–10, 19).

### Data availability

The GISAID datasets used in this study are available from the GISAID database (https://www.gisaid.org; EPI-SET-ID: EPI_SET_240925cs). The supplemental tables for the GISAID datasets are available in our GitHub repository (https://github.com/TheSatoLab/JN.1_full).

### Quantification and statistical analysis

Statistical significance was tested by one-way ANOVA with Tukey’s multiple comparisons test using GraphPad Prism 10 (GraphPad Software, MA, USA) unless otherwise noted. The values p < 0.05 were considered statistically significant (∗*p* < 0.05, ∗∗*p* < 0.01, ∗∗∗*p* < 0.001, ∗∗∗∗*p* < 0.0001). In the time-course experiments (Fig. 4A), a multiple regression analysis including experimental conditions as explanatory variables and timepoints as qualitative control variables was performed to evaluate the difference between experimental conditions across all timepoints. The initial timepoint was removed from the analysis. The *P* value was calculated by a two-sided Wald test. Subsequently, familywise error rates (FWERs) were calculated by the Holm method. These analyses were performed in R v4.1.2 (https://www.r-project.org/). All assays were performed independently at least 2 times.

## CONSORTIA

The Genotype to Phenotype Japan (G2P-Japan) Consortium: Hirofumi Sawa, Kenji Shishido, Hayayo Ito, Yu Kaku, Naoko Misawa, Arnon Plianchaisuk, Ziyi Guo, Alfredo Hinay Jr., Keiya Uriu, Yusuke Kosugi, Shigeru Fujita, Jarel M. Tolentino, Luo Chen, Lin Pan, Mai Suganami, Mika Chiba, Kyoko Yasuda, Keiko Iida, Naomi Ohsumi, Kazuhisa Yoshimura, Kenji Sadamasu, Mami Nagashima, Hiroyuki Asakura, Isao Yoshida, So Nakagawa, Akifumi Takaori-Kondo, Kotaro Shirakawa, Kayoko Nagata, Ryosuke Nomura, Yoshihito Horisawa, Yusuke Tashiro, Yugo Kawai, Sayaka Deguchi, Yukio Watanabe, Ayaka Sakamoto, Naoko Yasuhara, Takao Hashiguchi, Tateki Suzuki, Kanako Kimura, Jiei Sasaki, Yukari Nakajima, Hisano Yajima, Takashi Irie, Ryoko Kawabata, Kaori Tabata, Terumasa Ikeda, Hesham Nasser, Begum MST Monira, Ryo Shimizu, Michael Jonathan, Yuka Mugita, Otowa Takahashi, Takamasa Ueno, Mako Toyoda, Akatsuki Saito, Maya Shofa, Yuki Shibatani, Tomoko Nishiuchi.

## ACKNOWLEDGMENTS

We would like to thank all members of The Genotype to Phenotype Japan (G2P-Japan) Consortium. We thank H. Kubo, M. Tetsuka, and S. Shimamura for their secretory work and M. Hanazaki, H. Murota, A. Shigeno, I. Kida, M. Kurion, and Z. Guo for their technical assistance. We gratefully acknowledge the numerous laboratories worldwide that have provided sequence data and metadata to GISAID. A full list of originating and submitting laboratories for the sequences used in our analysis can be found at https://www.gisaid.org using the EPI-SET-ID: EPI_SET_240925cs.

This study was supported in part by AMED SCARDA Japan Initiative for World-leading Vaccine Research and Development Centers “UTOPIA” (JP223fa627001, to K.S.), AMED SCARDA Program on R&D of new generation vaccine including new modality application (JP223fa727002 to K.S.); AMED SCARDA Hokkaido University Institute for Vaccine Research and Development (HU-IVReD) (223fa627005 to T.F., and K.M.); AMED Project for Advanced Drug Discovery and Development (JP21nf0101627 to T.F.); AMED Research Program on Emerging and Re-emerging Infectious Diseases (JP21fk0108493, JP22fk0108617, JP22fk0108516 to T.F.; JP22fk0108146, to K.S.; JP21fk0108494 to G2P-Japan Consortium, K.M., S.T., T.F., and K.S.); AMED Research Program on HIV/AIDS (JP22fk0410039 to K.S.); AMED Japan Program for Infectious Diseases Research and Infrastructure (JP22wm0125008, to K.M.); AMED CREST (JP21gm1610005 to K.T.; JP22gm1610008 to T.F.); JST PRESTO (JPMJPR22R1 to J.I.); JST CREST (JPMJCR20H4 to K.S.); JSPS KAKENHI Fund for the Promotion of Joint International Research (International Leading Research) (JP23K20041 to K.M., K.S., and T.F.); JSPS KAKENHI Grant-in-Aid for Scientific Research B (21H02736 to T.F.); JSPS KAKENHI Grant-in-Aid for Early-Career Scientists (20K15767 to J.I.); JSPS Core-to-Core Program (Advanced Research Networks) (JPJSCCA20190008 to K.S.); World-leading Innovative and Smart Education (WISE) Program 1801 from the Ministry of Education, Culture, Sports, Science and Technology (MEXT) (to N.N.); Ministry of Health, Labour and Welfare (MHLW) under grant 23HA2010 (to N.N., and K.M.); The Cooperative Research Program (Joint Usage/Research Center program) of Institute for Life and Medical Sciences, Kyoto University (to K.S.); the Joint Research Program of Institute for Genetic Medicine, Hokkaido University (to K.Y., and T.T.) Akiyama Life Science Foundation (to T.T.); Japan Antibiotics Research Association (to T.T.); Hirose Foundation (to T.T.); The Tokyo Biochemical Research Foundation (to K.S.); Takeda Science Foundation (to T.F.); Hokkaido University Support Program for Frontier Research (to T.F.); and Tsuchiya Mitsubishi Foundation (to K.S.).

## AUTHOR CONTRIBUTIONS

S.T., J.I., K.S., K.M., S.T., T.T., and T.F. designed the experiments. S.T., T.M., N.N., K.O., L.W., Y.O., Y.M., J.L., R.H., Y.M., K.Y., M.N., J.I., K.T., K.M., and T.T. performed the experiments. S.T., and N.N. analyzed the data. S.T., T.T., and T.F. wrote the manuscript.

## Supplementary files

**Fig. S1.**
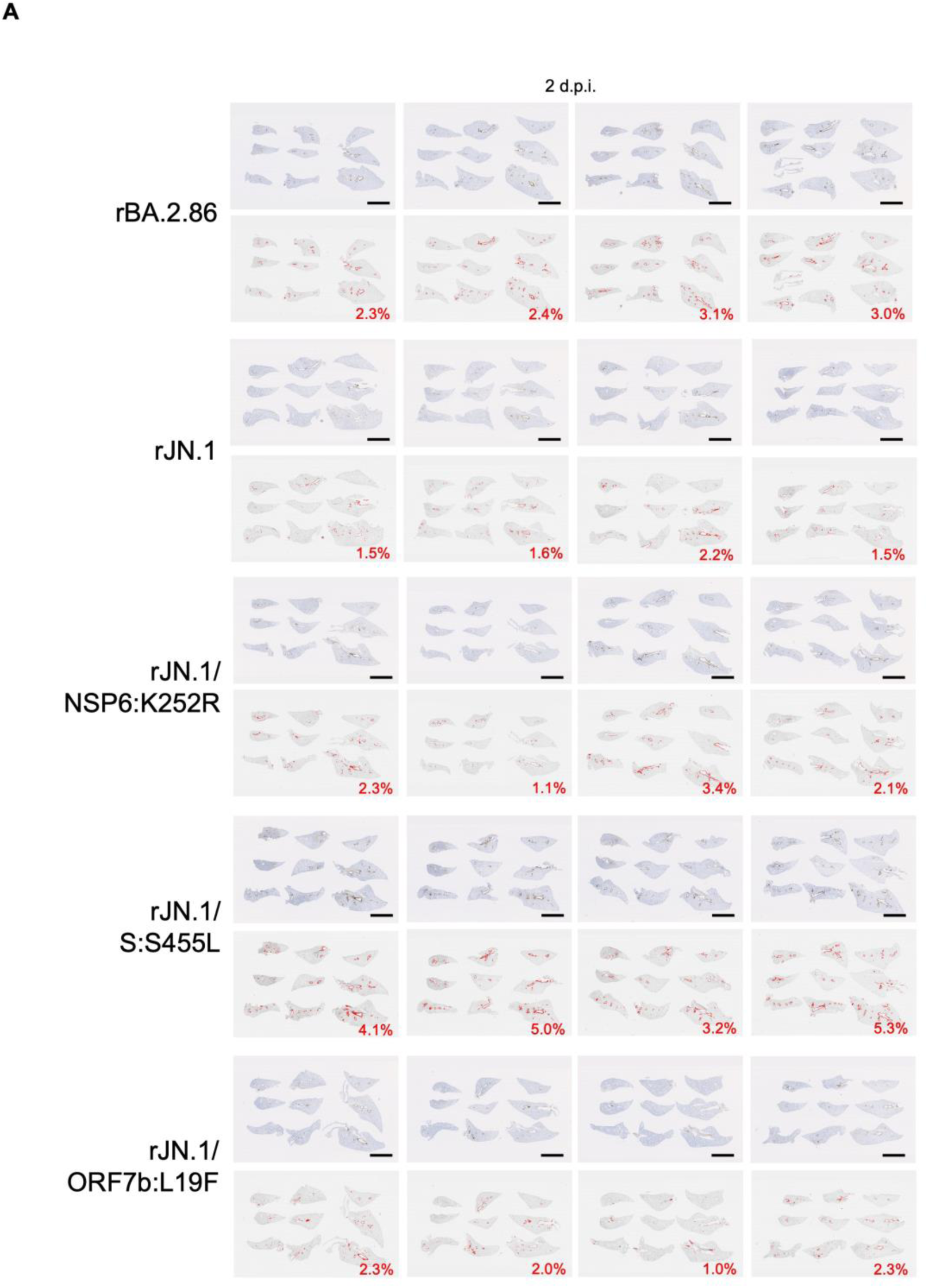
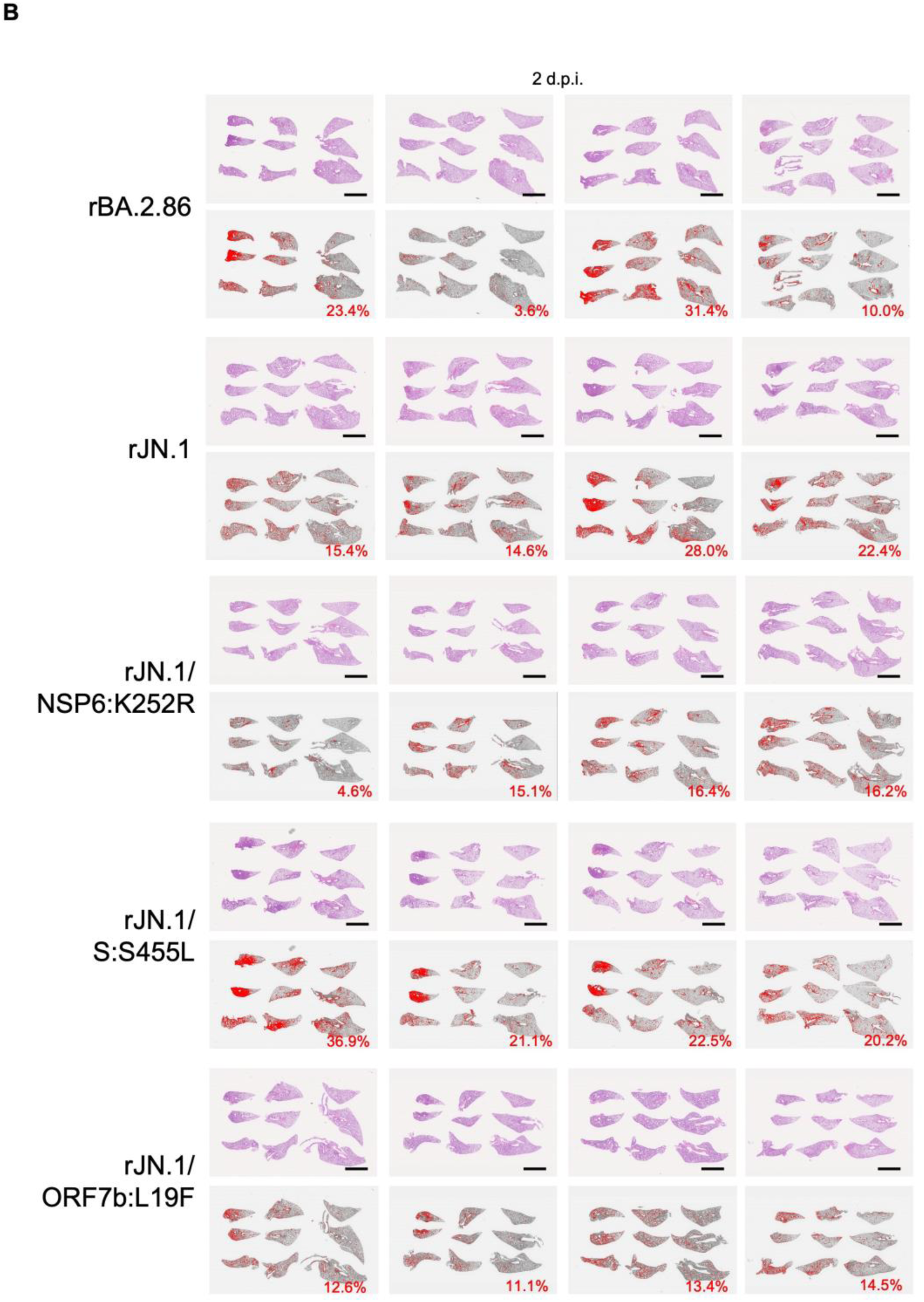
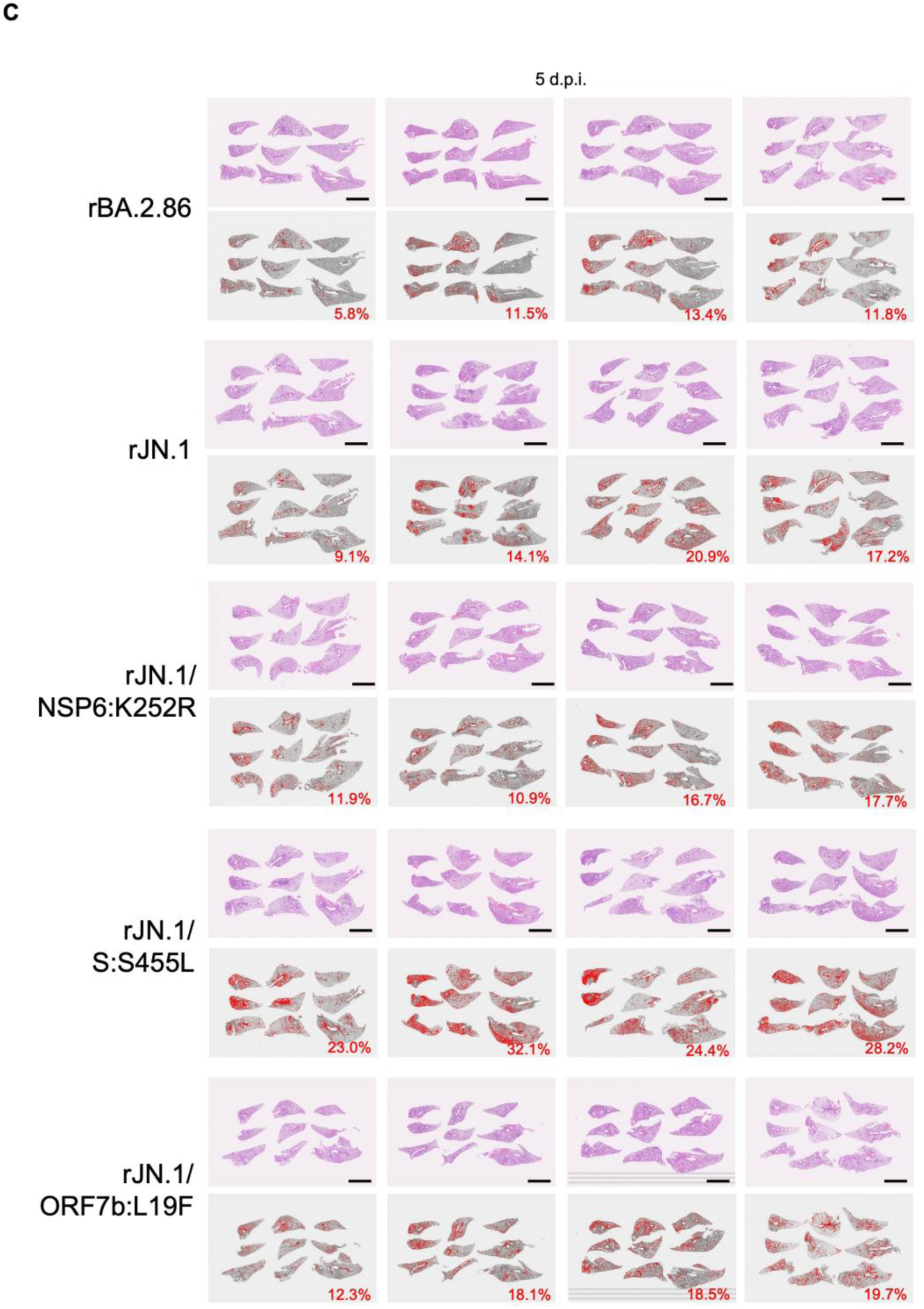
Pathological features of the infected hamsters, related to Figures 4C and 4D. **(A)** Immunohistochemistry of the viral N protein in the lungs of infected hamsters at 2 d.p.i. The percentages of N-positive cells in whole lung lobes are shown in the lower panels for each infection. Scale bars, 5 mm. **(B and C)** Hematoxylin and eosin staining of the lungs of infected hamsters at 2 d.p.i. **(B)** and 5 d.p.i. **(C)** Inflammatory areas with type II pneumocytes are shown in red and the percentage calculated in the lower panels for each infection. Scale bars, 5 mm.

